# Plasticity of sex-biased aggression in response to the sex of territory intruders in an African cichlid fish, *Julidochromis marlieri*

**DOI:** 10.1101/2024.05.22.595328

**Authors:** Ry Dennis, Kelsey J. Wood, Suzy C.P. Renn, Andrew P. Anderson

## Abstract

Behavior is often linked to gonadal sex; however, ecological or social environments can induce plasticity in sex-biased behaviors. In biparental species, pairs may divide offspring care into two parental roles, in which one parent specializes in territory defense and the other in nest care. The African cichlid fish *Julidochromis marlieri* displays plasticity in sex-biased behaviors. In Lake Tanganyika, *J. marlieri* form female-larger pairs in which the female is more aggressive than the male who performs more nest care, but under laboratory conditions, male-larger pairs can be formed in which these sex-biased behaviors are reversed. We investigated the influence of social environment on behavior by observing how individuals in both pair-types respond to conspecific intruders of either sex. We examined behavioral responses to three factors: sex of the subject, relative size of the subject, and the sex of the intruder. We confirm that relative size is a factor in behavior. The larger fish in the pair is more aggressive than the smaller fish is towards an intruder. While neither fish in the female-larger pairs varied their behaviors in response to the sex of the intruder, both members of the male-larger pairs were sensitive to intruder sex. Both individuals in the male-larger pairs engaged in more biting behaviors towards the intruder. Intruder biting behaviors strongly correlated with the biting behavior of the larger individual in the pair and occurred more frequently when encountering pairs with same sex as the larger fish when compared to pairs with the same sex as the smaller fish. Our results support the role of the social environment as a contributor in the expression of sex-biased behavior.

## Introduction

In many animals, there are sex-biased traits that can range from sex-specific to near monomorphic and the degree to which those traits are heteromorphic can change depending on social and environmental contexts. Aggression, parental care, and territory defense are behaviors that are often sex-biased to varying degrees (Huntingford and Turner 1987; Boesch 1992). Although females and males are both capable of performing a suite of behaviors, in bi-parental species task partitioning is often an adaptive solution in which parental roles can be divided by sex, size, or morphotype (convict cichlids: Snekser and Itzkowitz 2014, cichlids: Erlandsson & Ribbink 1997, White-Throated Sparrow: Tuttle 2003). While sexually heteromorphic behaviors can map strictly to gonadal sex, many species show plasticity for sex-biased behaviors. The degree to which each individual expresses a sex-biased behavior can be greatly influenced by biotic and abiotic environmental factors, including the social environment.

Sex-biased courtship behaviors have been repeatedly shown to plastically respond to environmental and social conditions. The intensity and direction of choosiness in both males and females as well as mating strategies can be influenced by food availability and population density (katydids and bushcrickets: Gwynne and Simmons 1990; Ritchie, Sunter, and Hockham 1998; locust: Pener and Yerushalmi 1998). With regard to the social environment, sex ratio can influence courtship roles in insects and fish species (butterfly *Acraea sp*.: Jiggins, Hurst, and Majerus 2000; two spotted goby: Forsgren et al.,2004; black striped pipefish: Silva et al., 2010) and parental care can shift in the absence of the typical caring sex or partner (burying beetles: Creighton et al., 2015; Suzuki and Nagano 2009; dendrobatid frogs: Ringler et al., 2015; strawberry poison-dart frog: Killius and Dugas 2014; dyeing poison frog: Fischer and O’Connell 2020). Complex factors in the social environment such as an intruder or audience can influence sex-biased behaviors (burying beetle: Ratz, Leissle, and Smiseth 2022; guppy: Plath et al., 2008; betta: Doutrelant, McGregor, and Oliveira 2001; Matos and McGregor 2002).

There is often a sex-bias in contest interactions which are also known to be influenced by size, with large size conferring an advantage in contests (cichlids: Barlow, Rogers, and Fraley 1986; Itzkowitz, Santangelo, and Richter 2001; Lehtonen et al., 2011; O’Connell and Hofmann 2012; Kidd et al., 2013). This effect and its interaction with sex has been studied in convict cichlids where males are generally larger than their mates and provide the majority of territory defense, whereas females are smaller and provide the majority of direct egg care. In experimentally size-reversed pairs, the degree of sex-biased behavior but not the direction is altered for both aggression and egg care (Itzkowitz et al., 2005).

Another group of cichlids, the African genus *Julidochromis* (tribe *Lamprologini*), offers a more extreme example of plasticity in which the direction of sex-bias can be reversed by manipulating the relative size of the animals in the pair. This tribe includes species with male-larger pairs in which males express more territorial behaviors (Taborsky and Limberger 1981) as well as species with female-larger pairs in which females express more territorial behaviors and males have smaller home ranges and spend more time at the nest (Ito, Yamaguchi, and Kutsukake 2017; Barlow 2005; Barlow & Lee 2005; Yamagishi and Kohda 1996; Kohda and Awata 2004).

For some of these species, the relative size pairing and the associated sex-biased behaviors have been shown to be plastic such that the larger fish is more aggressive regardless of sex and is more likely to take a second mate (Kohda and Awata 2004; Wood et al., 2014; Awata et al., 2006; Ito, Yamaguchi, & Kutsukake 2017; Yamagishi and Kohda 1996). These studies describe a system in which the size-mediated, sex-biased plasticity can reverse which individual in the pair performs the majority of one behavior or another.

Sex biased plasticity in territory defense was demonstrated for *J. marlieri* for the two pairing types, female-larger and experimentally reversed male-larger, both while eggs were present in the nest and prior to a broodcare phase (Wood et al., 2014). In that study, the use of a heterospecific intruder precluded the investigation of complex social dynamics involving the interaction with an intruder of the same species, which is not only a potential territory threat but, depending on the sex of the intruder, also represents a potential mate and threat to the pair-bond. Alternating the sex of a conspecific intruder is necessary to understand the interaction of intruder sex and pair-type.

Here, we aim to address how plasticity of sex-biased behavior interacts with the sex of a conspecific intruder. We test the plastic sex-biased behavior of aggression in *Julidochromis marlieri*, a biparental cichlid that normally forms pairs with a female that is larger the male, by manipulating the social environment based on relative size, sex, and intruder sex. We do this by presenting different sex-larger pairs with sequential intruder challenges of different sexes.

Based on past research, we predict the larger fish in the pair, regardless of sex, will show more aggression toward the intruder, and that intrasexual aggression will be greater than intersexual aggression because the intruder would represent a threat to the pair-bond.

## Methods

### The Study Population

The *J. marlieri* used in this study were obtained from the hobby trade and maintained in circulating water under conditions to mimic Lake Tanganyika (630-650 µS/cm at pH 8.3, 28 ± 0.3°C) on a 12/12 light/dark cycle with 30 minutes of dusk and dawn and fed flake food to satiation. Tanks included gravel and terracotta tile nests. It is not possible to know how many generations these fish were removed from wild, their exact age, nor the genetic relatedness among individuals, yet breeding captive pairs perform the same types of behaviors as observed in the wild. The advantages afforded by the common garden environment, the ability to control relative size, the ease of observation allowed us to quantify the effect of social environment on pair behavior. This research adhered to the ASAB/ABS Guidelines for the Use of Animals in Research. All experiments were performed in accordance with relevant institutional and national guidelines for the care and use of laboratory animals reviewed and approved by the Institutional Animal Care and Use Committee of Reed College (protocol #1032007).

### Establishing Pairs

Two mixed sex population tanks (110 L 5-20 fish) were established, one with size bias for larger males and the other with size bias for larger females. When two fish in the community tank displayed pairing behaviors such as nest defense (Wood et al., 2014), they were designated as a pair and were weighed, measured, and had sex confirmed visually (Table 1). Pairs were then moved into the pair compartment of an observation tank (110 L) (Fig. 1) that was divided by an opaque acrylic divider, and further divided by a clear acrylic divider with small perforations for the intruder compartment. Five female larger pairs and three male larger pairs were successfully established and remained paired throughout the experiment. With regard to the STRANGE framework (Webster and Rutz, 2020), the need for voluntary pairing in the reverse size relationship may introduce bias in that not all individuals in the species may behave this way. All pairs were acclimated for three to four days. The same individuals remained paired throughout the experiment. Intruders, chosen from a different tank than those used to form pairings, were sexed, measured (Table 1) and housed individually.

**Figure 1:**
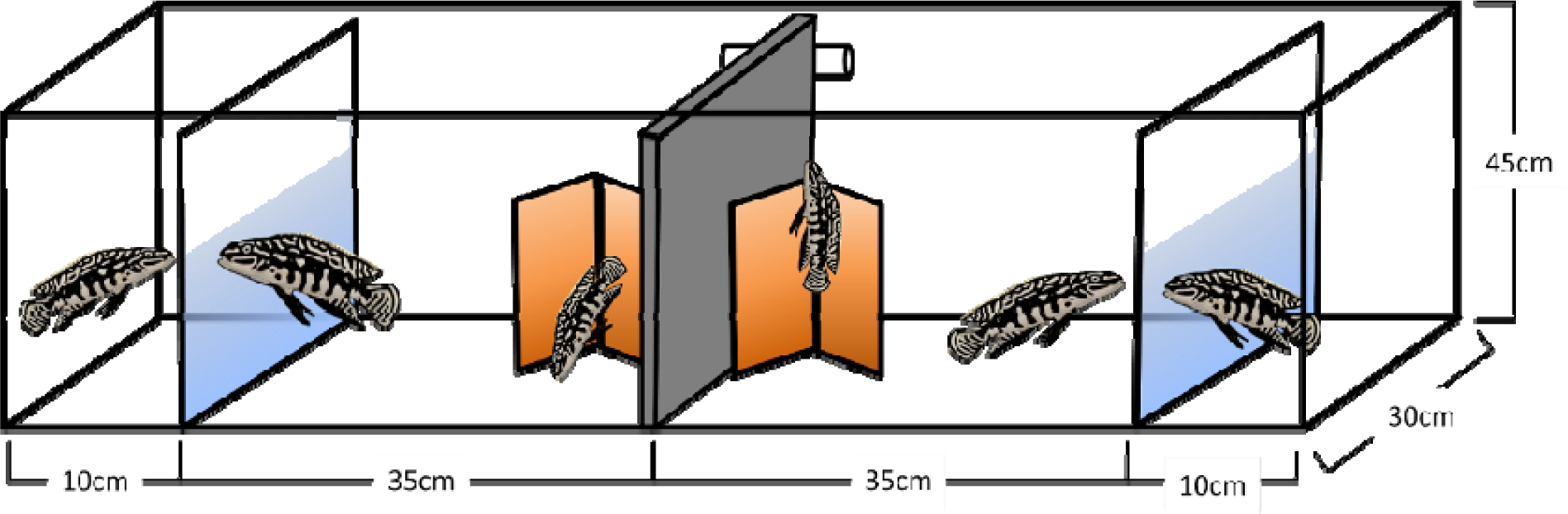
Observation tank with two pairs in the two pair compartments separated by an inserted opaque divider with an intruder in each intruder compartment separated from a pair by perforated clear dividers.

**Table 1:** Standard length (SL) and weight of Females (F) and Males (M) in female-larger (FL) and male-larger (ML) pairs. Intruder ID listed under the trial number, and the letter denotes female (F) or male (M) followed by the weight of the individual to demonstrate relative size of either sex intruder being sometimes larger and sometimes smaller than the smaller individual in the focal pair.

| Pair ID | SL (mm) |  | Weight (g) |  | Intruder Trials |  |  |  |
| --- | --- | --- | --- | --- | --- | --- | --- | --- |
|  | F | M | F | M | 1 | 2 | 3 | 4 |
| FL1 | 81 | 50 | 10.2 | 2.8 | F59 | M39 | F35 | M60 |
| FL2 | 83 | 50 | 10.2 | 2.4 | M63 | F59 | F39b | M37 |
| FL3 | 76 | 50 | 7.4 | 2.3 | F39a | M53 | F50 | M34 |
| FL4 | 70 | 53 | 5.9 | 2.4 | M50 | M37 | F39b | F54 |
| FL5 | 67 | 51 | 5.6 | 2.4 | M39 | F54 | F35 | M59 |
| ML1 | 42 | 81 | 1.6 | 8.6 | M60 | F59 | F39b | NA |
| ML2 | 45 | 68 | 1.8 | 5.1 | M37 | F39b | M53 | F50 |
| ML3 | 49 | 66 | 1.8 | 5.3 | M34 | F50 | F39a | M53 |

### Behavioral Observation

After acclimation, each pair was challenged with an intruder at about five hours after artificial sunrise on days 0, 2, 4, and 6, twice with female intruders and twice with male intruders, in a systematically varied order (Table 1). Video recording with a FujiFilm FinePix S8400W digital camera commenced as the intruder was placed in the small compartment and continued for 10 minutes, after which the intruder was removed. Behaviors were scored using BORIS (Friard and Gamba, 2016) by an observer blind to sex of all fish.

### Ethogram

We scored behaviors using an ethogram with five behaviors. “Out-of-nest” was measured as the time that each fish spent with no portion of its body or fin within the nest. “Close-to-divider,” a subset of out-of-nest, was measured as time that each fish spent within one body length of the divider and indicates overall interest in the intruder. “Bite” was scored as the number of times the fish bit or contacted the divider face-first regardless of opponent proximity on the other side of the divider. This same behavior was also scored for the intruders. Given the restricted space and lack of nest this was the only behavior scored for intruders In addition to these behaviors related to intruder inspection, we quantified two other social behaviors. Lateral-roll was scored as the number of times the subject’s body rotated along the anteroposterior axis such that the dorsal and ventral axis became roughly parallel with the tank floor. This behavior was not mutually exclusive with any other behavior states. “Bite mate” was scored when either member of the pair swam rapidly towards its mate or opened and closed its mouth while within 1 body length of its mate.

### Statistical Analysis

Data were processed and statistical analysis performed in R version 4.2.1 (R Core Team 2017) using tidyverse (Wickham et al., 2019), lme4 (Bates et al., 2015), and multcomp (Hothorn, Bretz, and Westfall 2008) packages. We applied general linearized mixed effects models (GLMM) with the behaviors of interest as the response variables. We investigated three factors: effects of relative size of the individual in the pair, sex of the individual, sex of the intruder, two-way interactions between each of the factors, and a three-way interaction between the three factors. Trial number was included as a fixed effect while intruder ID and individual ID were included as random effects. This model was chosen based on Akaike information criterion (AIC) (Akaike 1974) as it performed the best for the majority of behaviors and did not dramatically compromise the others. The behaviors bite, close-to-divider, and lateral-roll (generally assumed to be a submissive behavior) were modeled with a Poisson distribution, while time out-of-nest was modeled with a Gaussian distribution after assessing the distribution of data for each behavior. *Post-hoc* pairwise comparisons (t-test) were run for 16 contrasts of interest. We compared subjects only when they had two of the three conditions (subject sex, subject size, intruder sex) in common or were in the same pair type facing the same intruder sex. Unadjusted P-values are reported and we provide the adjusted alpha value for Bonferroni correction (P = 0.003125) and indicate when P-values are below this threshold for the 16 *post-hoc* contrasts.

Due to infrequent occurrence of bite-mate, this behavior was not analyzed with the GLMM. Instead, we summed the total occurrences of bite-mate for both intruder sexes thus trial number and intruder ID could not be included in a model thus this behavior. Furthermore, since only large individuals performed bite-mate subject size could not be included, thus this behavior was analyzed with a simple two-way ANOVA using only individual sex and intruder sex as factors.

To analyze the behavior of the intruder a different GLMM was required because the opponent ID was not recorded (we did not infer directed intent). We used pair-type and the intruder’s sex as categorical variables. To address social interaction from the perspective of intruder, we included the number of bites by the larger fish in the model since the larger fish engaged in more biting behavior. We used intruder ID and pair ID as the fixed effects with a Poisson distribution. *Post-hoc* pairwise comparisons (t-test) were run for four contrasts of interest. Unadjusted P-values are reported and we provide the adjusted alpha value for Bonferroni correction (P = 0.0125) and indicate when P-values are below this threshold for the four *post-hoc* contrasts.

## Results

### Pairing Success

Seven pairs remained bonded throughout all four intruder trials (5 species-typical female-larger and 2 experimentally-reversed male-larger pairs), but we included one additional male-larger pair that split between trials 3 and 4.

### Trial number effect

There was a general pattern for pairs to perform more behaviors upon subsequent trials (Fig. 2). The increased behavior was most pronounced for bites and close-to-divider (subsequent trials relative to the first: p < 0.001) but was also significantly increased for lateral-roll in trials 2 and 4, relative to trial 1 (p < 0.05).

**Figure 2:**
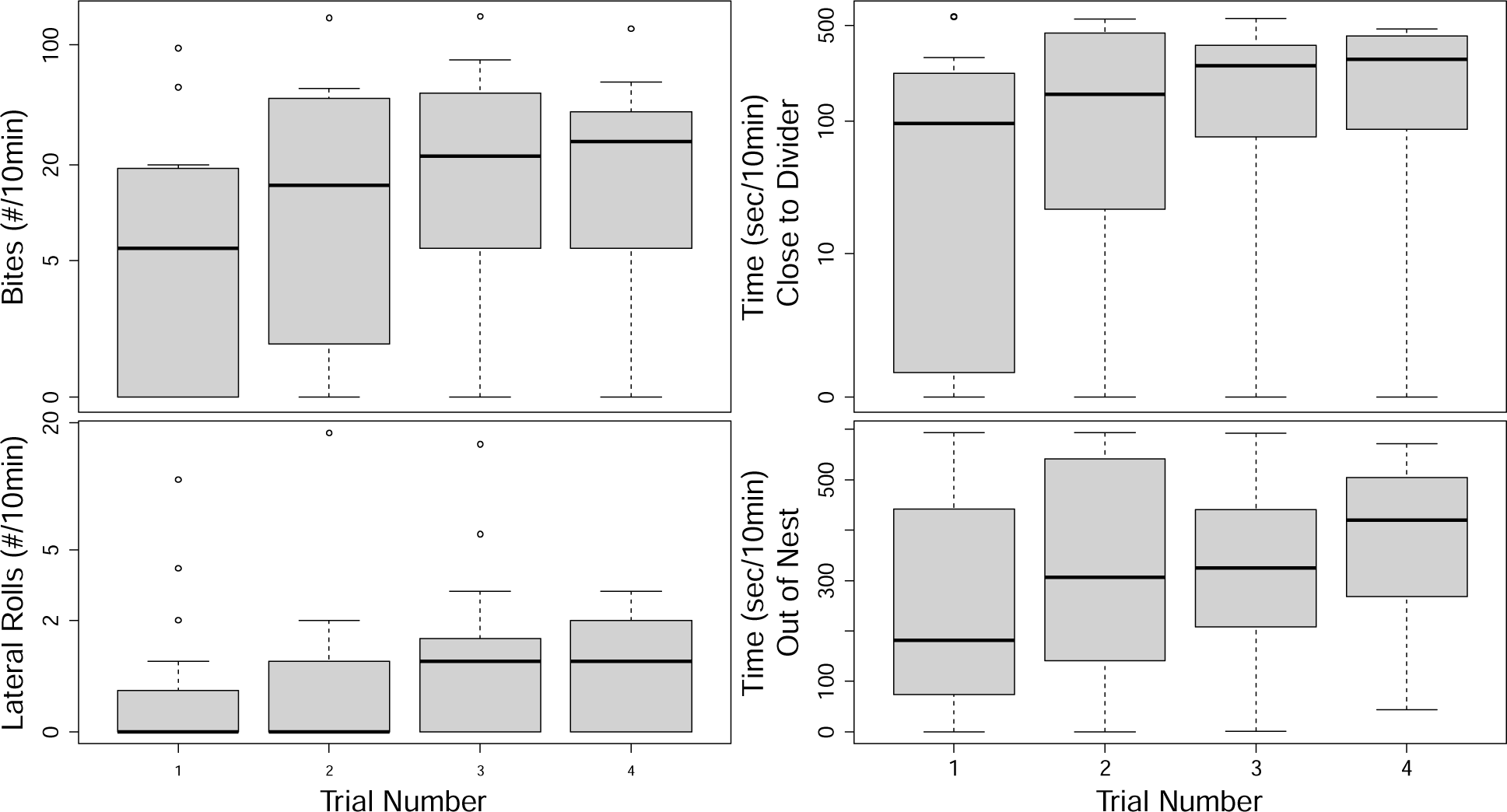
Boxplot showing total events or time in seconds for all focal fish in each behavioral trial.

### Bite

In order to quantify bites directed at the intruder, we counted the number of times each fish in the focal pair struck or bit at the transparent divider. As predicted, relative size within the pair (larger or smaller) had a significant effect on bites (p = 0.016) (Table 2), with the larger fish performing more bites (Fig. 3a). Neither sex of the subject nor sex of the intruder had significant main effects on the number of bites (Table 2). However, there were significant two-way interactions between the relative size of the subject and the sex of the intruder (p < 0.001), as well as between sex of the subject and the sex of the intruder (p < 0.001). There was also a significant three-way interaction between relative size, sex of the subject, and sex of the intruder (p < 0.001) (Table 2).

**Figure 3:**
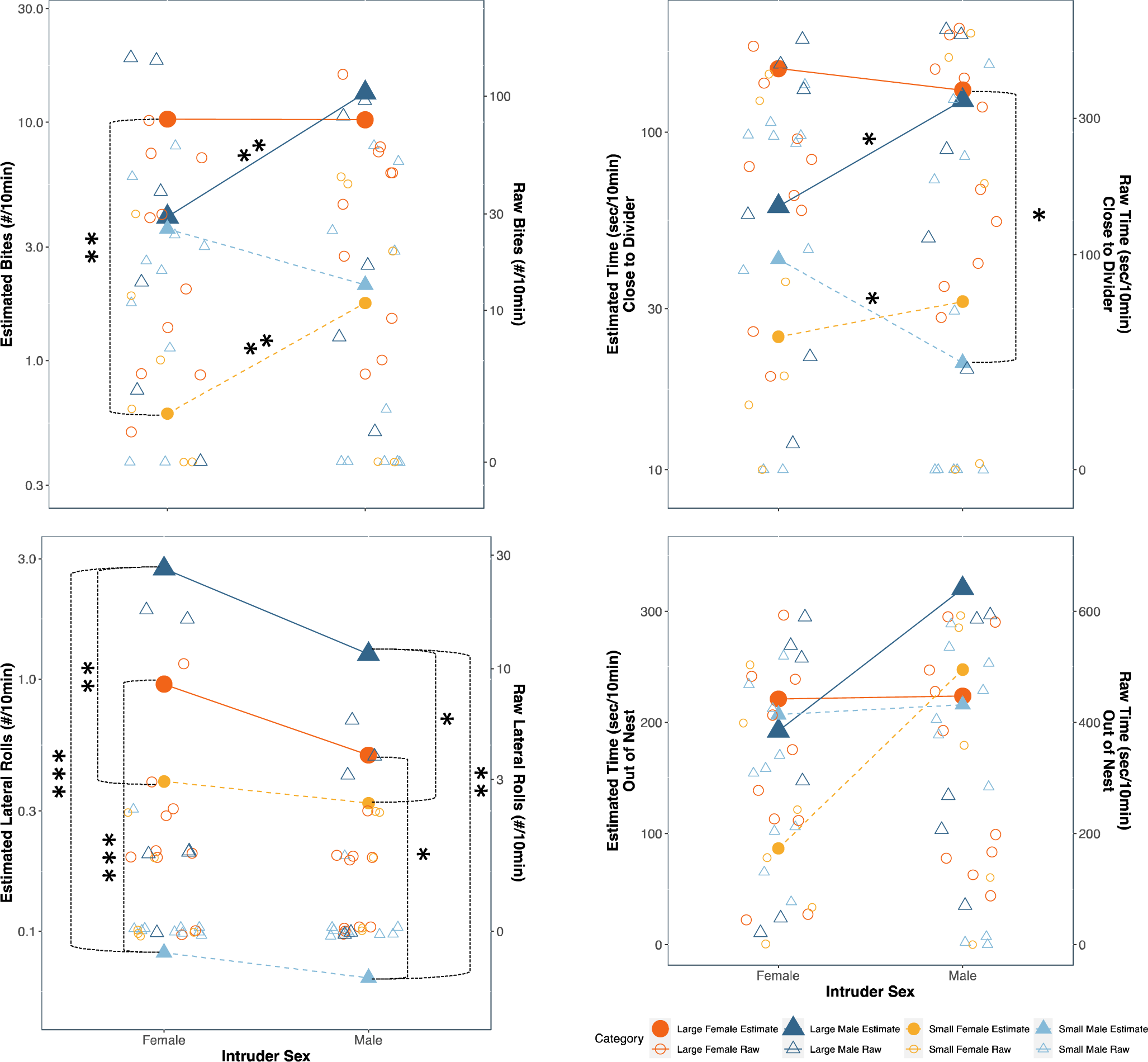
Reaction Norm Plot for the linear estimate for behavioral measures over the course of 10 minutes in response to female intruders and male intruders for fish in species-typical female-larger pairs (large females=large red circles; small males=small light blue triangles) and fish in experimentally-reversed male-larger pairs (large males=large dark blue triangles and small females=small orange circles). Open shapes represent raw values of the behavioral measure prior to adjustment from multi-variate model (note different y-axis values). Asterisks indicate level of significance for pairwise comparisons (*—p < 0.10, **—p < 0.05, ***—p < 0.003125 Bonferroni correction).

**Table 2:**
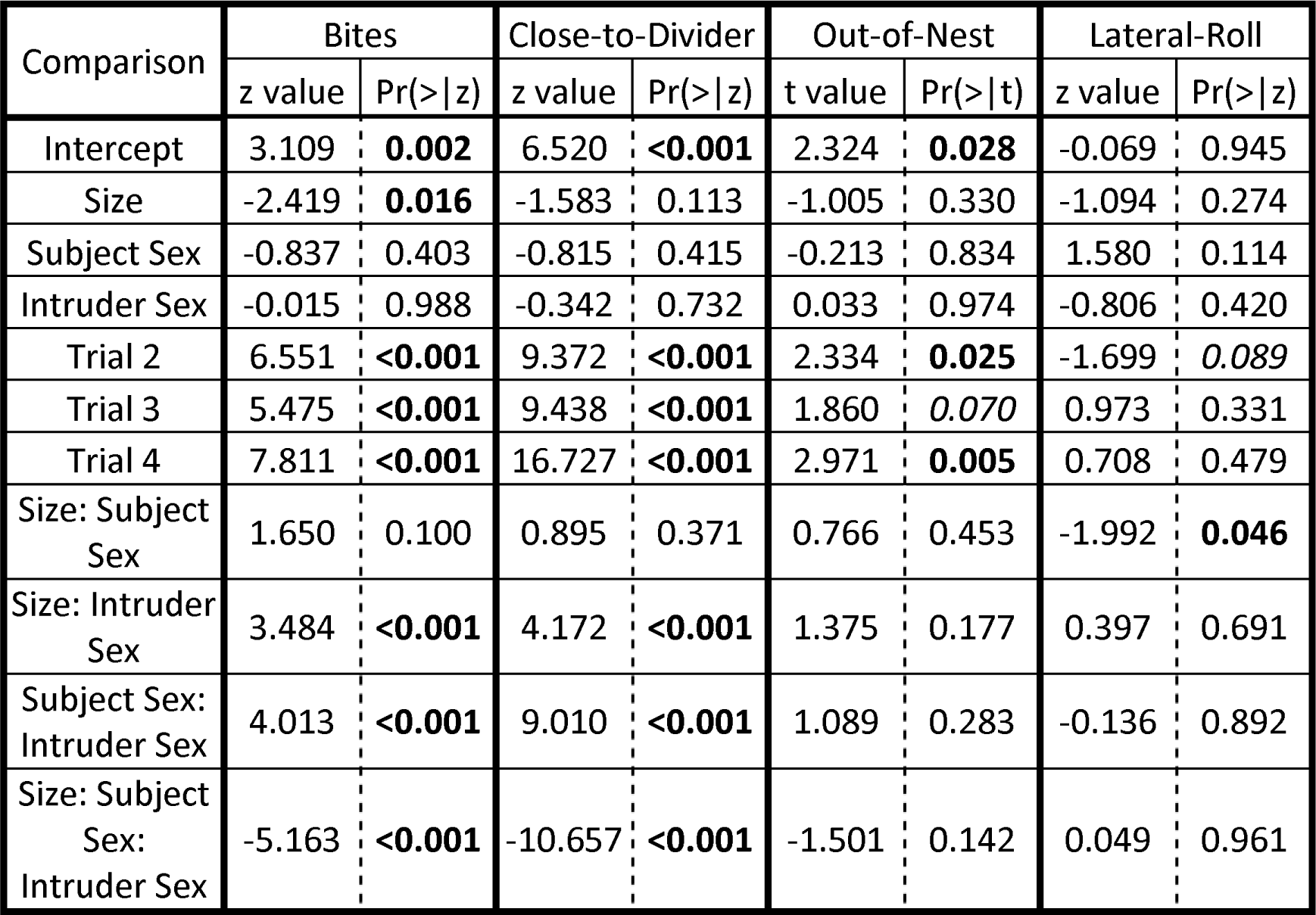
GLMM results for fixed and interaction effects. Z values, for Poisson distributed data, and t values, for normally distributed data, are shown to the left and p values are shown to the right. Bold values represent statistically significant effects at alpha = 0.05 and *italics* represent trends at alpha = 0.1.

*Post-hoc* pairwise comparisons show relationships between conditions. In species-typical female-larger pairs, the sex of the intruder did not have an effect on the number of bites performed by either fish in the pair (large females: p = 0.988, small males: p = 0.196) (Table 3). However, in experimentally-reversed male-larger pairs, the males had significantly more bites toward male intruders than toward female intruders (p=0.008), as did their smaller female mate (p=0.022) (Fig. 3a). Larger females in species-typical pairs bite more than the smaller females in experimentally-reversed pairs when presented with a female intruder (p=0.016) (Fig. 3a). While none of the P-values are significant following Bonferroni correction, they point to the underlying patterns of the significant factors found in the linear model.

**Table 3:** Pairwise comparisons results. Under the behavior measured, Chi-squared values are shown to the left and p values are shown to the right. **Bold** values represent statistically significant comparisons at alpha = 0.05 and *italics* represent trends at alpha = 0.1. When using Bonferroni correction for multiple tests (n =16) the new significant threshold for alpha = 0.003125. The leftmost column shows the pairwise comparison being performed and the second and third columns show the condition of the comparison.

| Compar-<br>ison | Conditions |  | Bites |  | Close- to-<br>Divider |  | Out-of-Nest |  | Lateral-Roll |  |
| --- | --- | --- | --- | --- | --- | --- | --- | --- | --- | --- |
|  |  |  | Chisq Pr(>\|Chisq\|) |  |  |  |  |  |  |  |
| Intruder<br>Sex<br>F vs. M | F<br>Larger | F Subject | <0.001 | 0.988 | 0.117 | 0.732 | 0.001 | 0.974 | 0.649 | 0.420 |
|  |  | M Subject | 1.673 | 0.196 | 2.733 | 0.098 | 1.537 | 0.215 | 0.026 | 0.872 |
|  | M<br>Larger | F Subject | 5.281 | **0.022** | 0.308 | 0.579 | 2.431 | 0.119 | 0.031 | 0.861 |
|  |  | M Subject | 7.018 | **0.008** | 2.790 | 0.095 | 0.013 | 0.909 | 0.683 | 0.409 |
| Subject<br>Sex<br>F vs. M | Large<br>Subject | F Intruder | 0.701 | 0.402 | 0.664 | 0.415 | 0.046 | 0.831 | 2.495 | 0.114 |
|  |  | M Intruder | 0.055 | 0.815 | 0.004 | 0.951 | 0.486 | 0.486 | 1.374 | 0.241 |
|  | Small<br>subject | F Intruder | 2.283 | 0.131 | 0.207 | 0.649 | 0.809 | 0.369 | 2.195 | 0.139 |
|  |  | M Intruder | 0.022 | 0.883 | 0.128 | 0.720 | 0.052 | 0.820 | 1.533 | 0.216 |
| Subject<br>Size<br>Large vs.<br>Small | F<br>Subject | F Intruder | 5.851 | **0.016** | 2.506 | 0.113 | 1.009 | 0.315 | 1.197 | 0.274 |
|  |  | M Intruder | 2.301 | 0.129 | 1.559 | 0.212 | 0.029 | 0.864 | 0.246 | 0.620 |
|  | M<br>Subject | F Intruder | 0.011 | 0.918 | 0.095 | 0.758 | 0.012 | 0.914 | 14.216 | **<0.001** |
|  |  | M Intruder | 2.691 | 0.101 | 2.366 | 0.124 | 0.567 | 0.451 | 5.462 | **0.019** |
| Between<br>mates<br>F vs. M | F<br>Larger | F Intruder | 1.178 | 0.278 | 1.656 | 0.198 | 0.016 | 0.901 | 8.540 | **0.003** |
|  |  | M Intruder | 2.669 | 0.102 | 3.394 | 0.065 | 0.005 | 0.945 | 2.999 | 0.083 |
|  | M<br>larger | F Intruder | 2.180 | 0.140 | 0.475 | 0.491 | 0.530 | 0.467 | 5.787 | **0.016** |
|  |  | M Intruder | 2.498 | 0.114 | 1.131 | 0.288 | 0.236 | 0.627 | 3.344 | 0.067 |

### Close-to-divider

As another measure of intruder interest, we quantified the time each fish in a focal pair spent close-to-divider, which could indicate inspection of the intruder. While there were no main effects of relative size of the subject, the sex of the subject, nor the sex of the intruder, there were significant two-way interactions between the relative size of the subject and the sex of the intruder (p < 0.001), between the sex of the subject and the sex of the intruder (p < 0.001), as well as a three-way interaction between relative size, sex of the subject, and sex of the intruder (p < 0.001) (Table 3).

Pairwise comparisons revealed differences according to the pairing types. In the species-typical female-larger pairs, the males tended to spend less time at the divider when presented with a male intruder than with a female intruder (p = 0.098). Conversely, in the experimentally-reversed male-larger pairs, the males tended to spend more time at the divider when presented with a male intruder than with a female intruder (p = 0.095) (Fig. 3b). As expected for species-typical female-larger pairs, the large females generally spent more time close-to-divider than their smaller male mates with a larger difference for male intruders (male intruders: p = 0.065) (Fig. 3b).

### Out-of-nest

As an additional measure, we quantified the time each fish in a focal pair spent with its body fully out of the nest, which could indicate vigilance, a trade-off against nest maintenance (Fig. 3c). While there were no significant main effects, two-way, or three-way interactions (Table 2), it is noteworthy that, similar to the behaviors described above, when presented with male intruders, the subjects in experimentally-reversed male-larger pairs did increase the time spent out-of-nest (large males: p = 0.215; small females: p = 0.119) but this was not the case for species-typical female-larger pairs (large females: p = 0.974; small males: p = 0.909) (Fig. 3c).

### Lateral-roll

We scored the relatively rare lateral-roll behavior (Fig. 3d). The only significant effect was a two-way interaction between the relative size of the subject and its sex (p = 0.046) (Table 3).

Pairwise comparisons showed that the larger subjects tended to perform more lateral-rolls than their smaller mates. In species-typical female-larger pairs, the females performed more lateral-rolls than their mates, although this difference was only significant when the intruder was female (female intruder: p = 0.003—below Bonferroni threshold; male intruder: p = 0.083). In experimentally-reversed male-larger pairs, the males performed more lateral-rolls than their mates and again the difference was only significant when the intruder was female (female intruder: p = 0.016; male intruder: p = 0.067). The large males in experimentally-reversed male-larger pairs increased the number of rolls performed compared to smaller males in species-typical pairs (female intruders: p < 0.001—below Bonferroni threshold; male intruder: p = 0.019) (Fig. 3d). By contrast, there was no significant difference in the number of lateral-rolls between larger females in species-typical pairs and smaller females in experimentally-reversed pairs (Table 3).

### Bite Mate

Because we considered it possible that the intruder would be perceived as a potential mate (*i.e.* threat to the pair bond), we attempted to determine how sex of the intruder would impact the pair dynamics in terms of bites directed at the mate. These bites were very rare (18 occurrences across 12 of the 31 total observations). Only large fish ever bit their mate. We ran a two-way ANOVA for sex of the subject and the sex of the intruder, looking only at the larger individuals, but there were no significant effects (subject sex: p = 0.511, intruder sex: p = 0.714, subject sex:intruder sex: p = 0.401).

### Intruder Bites

To address the role of the intruder’s behavior, we scored intruder bites and found that the number of bites made by the larger fish in the pair is a strong predictor of the intruder’s bite behavior (p < 0.0001) consistent with the idea that the intruders are responding to the pair. The interaction of pair type and intruder sex was highly significant the (p < 0.0001) such that intruders showed the greatest amount of biting behavior when they were the same sex as the larger fish in the pair. Independently, both the pair type, with female larger pairs eliciting more intruder biting the male larger pair (p = 0.0457) and intruder sex with females exhibiting more biting than males (p = 0.0199), were also significant (Fig 4).

**Figure 4.**
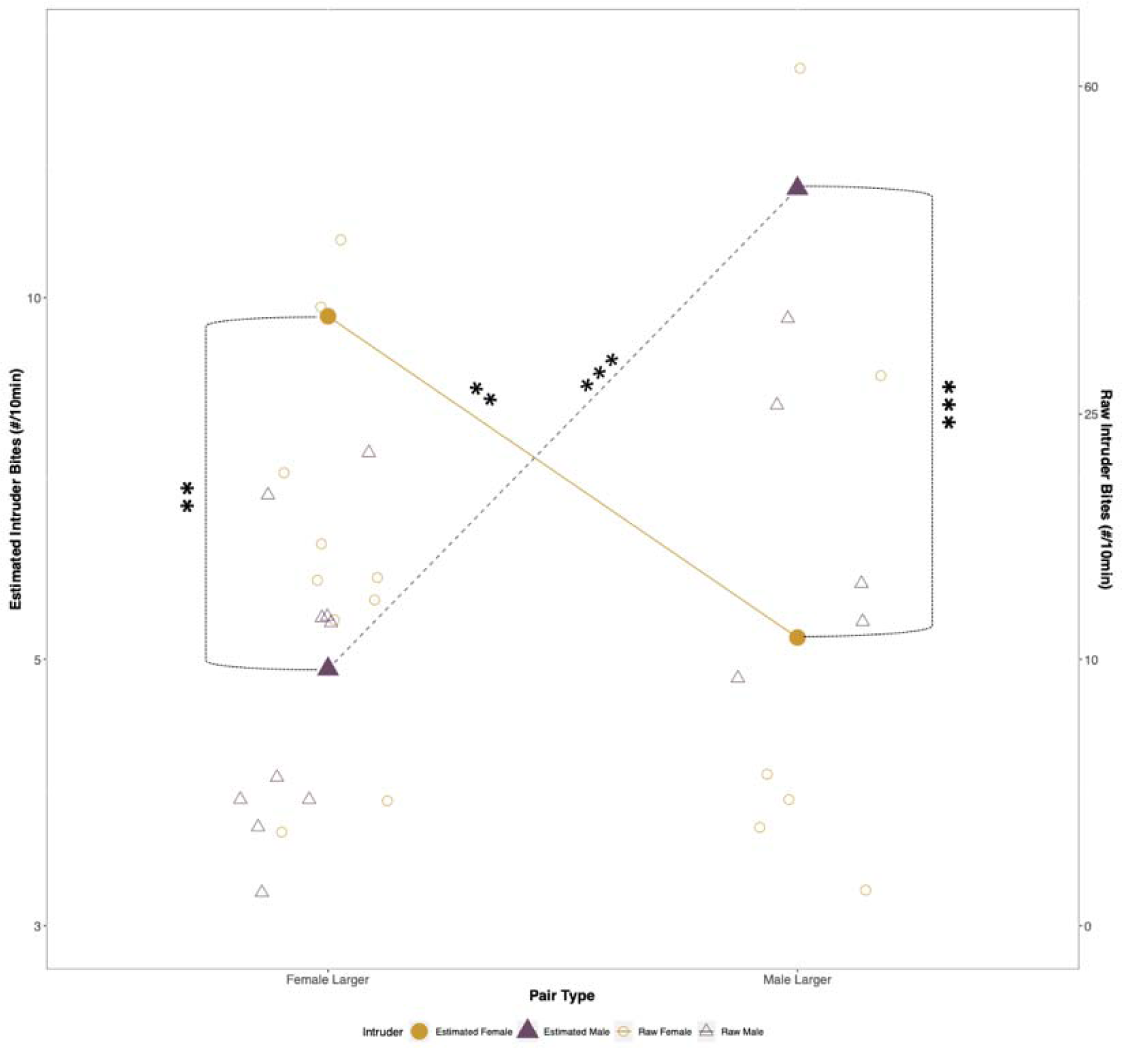
Reaction Norm Plot for the linear estimate for number of bites done by the intruding fish over the course of 10 minutes by sex (orange circle = female intruder, green triangle = male intruder) in response to species-typical female-larger pairs (left side) and fish in experimentally-reversed male-larger pairs (right side). Open shapes represent raw values of the behavioral measure prior to adjustment from multi-variate model (note different y-axis values). Asterisks indicate level of significance for pairwise comparisons (*—p < 0.10, **—p < 0.05, ***—p < 0.0125 Bonferroni correction).

## Discussion

Our results confirm that behavior in *J. marlieri* is not determined by the sex of subjects but rather is influenced by the social environment in terms of the relative size of the individuals in the pair (*J. marlieri*: Wood et al., 2014). While previous work reported a reversal in sex-bias for aggression and nest-care related behaviors, those studies did not manipulate additional variables of the social environment. Here, we also demonstrate that interest in an intruder is modulated by the interactions between the relative size of the subject, the sex of the subject, and the sex of the intruder. This pattern was strong and evident in multiple behaviors related to aggression, territory defense, and vigilance, even though not all differences were statistically significant. The clear pattern revealed that the larger fish in the pair is more attentive to the intruder than the smaller fish regardless of pairing type (male-larger vs. female-larger), but for the experimentally-reversed male-larger pairs, behavior was also influenced by the sex of the intruder.

We had hypothesized that the larger aggressive fish in both pairing types would vary their behavior based on the sex of the intruder, increasing intrasexual interactions and decreasing intersexual interactions as has been shown in other species (Yellow-Breasted Chats: Mays and Hopper 2004; review: Pandolfi, Scaia, and Fernandez 2021). Relatively larger males did show the expected higher intrasexual biting behavior and attentiveness towards the intruder (close-to-divider measure); however, this pattern was not seen for smaller female subjects, and neither sex in the species-typical female-larger pairings significantly varied biting in response to the sex of the intruder. A difference in responsiveness to the sex of the intruder has been seen in Tibetan Ground Tits in which the males (species-typical territory holders) exhibit high levels of aggression regardless of the intruder’s sex while the females show a reproducible plastic response (Guo et al., 2020). The pattern we observed suggests differences in priorities for females and males depending on their relative size within the pair and resulting social role within the pair. As discussed below, we propose two possible explanations: 1) experimentally-reversed male-larger pairs are more sensitive to the sex of an intruder because these pairs are less stable than the species-typical female-larger pairings, or 2) the attentive behavior performed by females toward male intruders actually represents courtship.

Stability of the pair could impact aggression against conspecific intruders which functions both as defense of a territory and defense of the pair bond, often including mate-guarding. The mere presence of a mate can promote mate-guarding (Meadow Pipit: Petrusková et al., 2007), but mate quality also plays a role in the level of aggression. Individuals may guard a higher quality mate more vigorously than they would a lower quality mate and conversely, a low quality mate may guard its mate more vigorously (review: Harts, Booksmythe, and Jennions 2016). Thus, the magnitude of intrasexual aggression is impacted by mate quality. For *J. marlieri* females, the species-typical, and therefore preferred, mate would be a relatively smaller male (Barlow and Lee 2005), but in our experimentally-reversed pairs, the females have pair-bonded with a larger, less preferred male. This atypical pairing may therefore expected to be less stable; neither member of the pair is with a mate of preferred size, and is therefore expected to show greater response to the sex of an intruder. The males in male-larger pairs exhibit a high level of attentiveness toward male intruders, who represent a territory threat as well as a threat to the pair-bond of the resident male, and they exhibit reduced attentiveness toward female intruders, who do not represent a threat to the pair-bond. Thus, larger males’ increased intrasexual interactions may represent an attempt to preserve their investment in their current mate by preventing the female’s access to preferred smaller males.

In experimentally-reversed pairs, the relatively smaller females showed overall low levels of interest in the intruder; however, there was a significant increase in intersexual bites directed at male intruders. This increase could be attributed to the small female taking cues from the large mate. Previous research in convict cichlids suggests that the smaller member in both species-typical and experimentally-reversed pair types follows the behavioral patterns set by the larger member (Itzkowitz et al., 2005). Alternatively, in experimentally-reversed male-larger pairings, the small female may see the intruder male as a potential mate. *Julidochromis* are primarily monogamous (Brichard 1989) but polyandry is reported for species with female-larger pairs (*J. ornatus:* Awata, Munehara, and Kohda 2005; Heg and Bachar 2006), so the small female is expected to show some interest in an additional mate. In *Julidochromis*, courtship often resembles aggression as both involve biting behaviors (Barlow and Lee 2005), and as a sex-role-reversed species, *J. marlieri* females could be expected to be the sex that performs courtship displays. Therefore, the behaviors observed in females may represent courtship both when the female is the smaller and when she is the larger fish in the pair.

As part of courtship, *J. marlieri* are known to bite and seemingly attack potential mates prior to pair bonding (Barlow and Lee 2005), and here we also see the larger fish in the pair engaged in more bites towards is partner. The inability for researchers to distinguish courtship signals from aggressive signals is a potential confounder in cichlid research (John et al., 2021). This ambiguity may explain the apparent lack of adjustment of the larger females’ biting behavior based on the sex of the intruder; she bites at male and female intruder equally. The female’s biting and hitting at the divider may be courtship when directed at male intruders. This idea is supported by the corresponding behavior of the intruder. Our data show that when the intruder experiences more bites from the larger fish they bite more, and when that larger fish is the same sex as the intruder the number of bites increases. Since same-sex intruders engage in an equally vigorous bite response this could be a sign of aggression by both individuals; whereas opposite-sex intruders do not respond to bites this may be interpreted as a similar courtship as seem within established pairs. This is predicted by female-biased courtship and male-biased choice in polyandrous species (review: Fritzsche et al., 2021). Even when the relative size-sex relationship is reversed, the direction of these sex-biases are often maintained (seahorses: Vincent 1994; two-spot goby: de Jong et al., 2009). While males play a role in the formation of a pair-bond, active courtship wouldn’t be expected to be part of male’s repertoire in a sex-role reversed species (Barlow and Lee 2005). Similar to aggression in convict cichlids (Itzkowitz et al., 2005), *J. marlieri* behavioral roles during pair-bond formation may change in magnitude but not in direction; thus, even as their relative size is reversed, females remain courters and males do not switch from chooser to courter. Our results support the hypothesis that the sex-bias of biting behaviors in *J. marlieri* can be reversed by changing the relative size of individuals in a pair, but behaviors related to pair-bond formation are not reversed by this social environmental factor.

The fish in our study were pair-bonded and territory holders as demonstrated by increased defense over the course of the experiment, a phenomenon that results from time investment in the territory and pair (midas cichlid: Barlow, Rogers, and Fraley 1986). *J. transcriptus* vary their responses to a challenger based on the outcome of recent contests (Hotta et al., 2021), duration since the last interaction (Hotta et al., 2014), and observations of previous interactions (Hotta et al., 2014; 2021). Characteristics of the interacting conspecific (*e.g.* intruders) could also influence sex-biased behaviors in *Julidochromis*. Subsequent interactions are also known to impact future investment in terms of a winner/loser effect (review: Hsu, Earley, and Wolf 2006), though this is thought to be weak in *Julidochromis* (Hotta et al., 2014; 2015; 2021). In order to uncover similarities and differences in courtship and aggression in this species, increased detail in the ethogram, prolonged observations, and additional social contexts are necessary. For example, the ambiguous lateral-roll behavior could signify aggression when combined with fin erection (Barlow and Lee, 2005), while signifying submission when accompanied by rapid retreat movement (Renn lab unpublished). Further research is needed to determine the significance of this behavior, but the current result suggests aggression as it is performed more often by the relatively larger fish. Examination of more complex social environments, such as the process of pair bond formation or interactions with neighboring territories, may reveal specific uses of these signals.

Here we have shown that paired *J. marlieri* adjust behavior in response to their relative size in the pair and the sex of conspecific intruders, and we suggest this represents a reversal in the sex-bias of territorial aggression while the species-typical female-biased courtship is retained. The plasticity of some behaviors should not be taken to indicate the plasticity of all behaviors. *Julidochromis* species present an excellent model for exploring the relative contributions of environmental factors toward the modulation of different sex-biased behaviors.

## AUTHOR CONTRIBUTIONS

**Dennis**: Investigation; methodology, data analysis and curation; writing – original draft. **Wood**: Conceptualization; methodology. **Renn**: Conceptualization; methodology; supervision; writing – review and editing. **Anderson**: Conceptualization; methodology; supervision; writing – review and editing; data analysis and curation.

## ACKNOWLEDGEMENTS

This work was supported by a REP supplement to NSF grant #1456486 and Reed College student research support through the Galakatos Science Research Fund. Lab members contributed to animal husbandry and critical discussion.

## DATA AVAILABILITY

The data used in this study and the code used to analyze them can be found at the github repository: https://github.com/AndersonDrew/JulidochromisIntruder

